# A fruitful endeavor: scent cues and echolocation behavior used by *Carollia castanea* to find fruit

**DOI:** 10.1101/532614

**Authors:** Leith B. Leiser-Miller, Zofia A. Kaliszewska, M. Elise Lauterbur, Brianna Mann, Jeffrey A. Riffell, Sharlene E. Santana

## Abstract

Frugivores have evolved sensory and behavioral adaptations that allow them to find ripe fruit effectively, but the relative importance of different senses in varying foraging scenarios is poorly known. Within Neotropical ecosystems, short-tailed fruit bats (*Carollia*: Phyllostomidae) are abundant nocturnal frugivores, relying primarily on plants of the genus *Piper* as a food resource. Previous research has demonstrated *Carollia* employ olfaction and echolocation to locate *Piper* fruit, but it is unknown how their sensory use and foraging decisions are influenced by the complex diversity of chemical cues that fruiting plants produce. Using wild *C. castanea* and their preferred food, *Piper scintillans*, we conducted behavioral experiments to test two main hypotheses: (1) foraging decisions in *C. castanea* are primarily driven by ripe fruit scent and secondarily by vegetation scent, and (2) *C. castanea* re-weight their sensory inputs to account for available environmental cues, such that bats rely more heavily on echolocation in the absence of adequate scent cues. Our results suggest that *C. castanea* requires olfactory information and relies almost exclusively on ripe fruit scent to make foraging attempts. Ripe fruit scent is chemically distinct from vegetation scent in *P. scintillans*, with a greater abundance of *β*-caryophyllene, germacrene D and *β*-elemene, and a few unique compounds. Although variation in echolocation call parameters was independent of scent cue presence, bats emitted longer and more frequent echolocation calls in trials where no fruit scent was present. Altogether, these results highlight the adaptations, plasticity, and potential constraints in the sensory system of neotropical fruit bats.

**SIGNIFICANCE STATEMENT:** Little is known about the relative importance of different senses and which plant cues are most important for fruit location by frugivores. We conducted behavioral experiments on short-tailed fruit bats (*Carollia castanea*), which use a combination of olfaction and echolocation to find ripe fruit, and their preferred food source (*Piper scintillans*) to test (1) which plant scent cues drive food selection and (2) if bats alter their echolocation behaviors based on which scent cues are present. We find that *C. castanea* rely almost exclusively on ripe fruit scent to forage, and echolocate more frequently when fruit scent is absent. Ripe fruit scent is chemically different from vegetation scent in *P. scintillans*, potentially providing a clear signal of food availability to mutualistic bats. These results highlight the sensory adaptations and behavioral flexibility of fruit bats as they navigate the cues provided by fruiting plants.

## INTRODUCTION

Animals rely on multiple sensory modalities to perform even the simplest ecological tasks (Burkhardt 1989; Siemers and Schnitzler 2000; Holland et al. 2005; Knaden and Graham 2016). One of the main goals of behavioral and sensory ecology lies in understanding the ability of different species to employ and modulate their sensory modes in the context of different environmental cues, and how the resulting behavioral decisions ultimately affect their ecology and evolution. Frugivores use various cues, including components of fruit color (Burkhardt 1989; Osorio & Vorobyev 2008, Melin et al. 2008, Hiramatsu et al. 2008; Valenta et al. 2013), shape (Helversen & Helversen 1998; Kalko & Condon 1998), and scent (Valenta et al. 2013, Sánchez et al. 2006) to find and select ripe fruit, and exhibit corresponding sensory specializations in their visual, auditory and/or olfactory systems to target those cues (Catania 1999; Muller et al. 2007; Vanderelst et al. 2010). Little is known, however, about what scenarios facilitate or constrain sensory system use and modulation during fruit location, selection, and acquisition in vertebrate frugivores. Insight into these processes would provide mechanistic understanding of the behaviors underlying their foraging ecology.

Frugivorous and omnivorous Neotropical leaf-nosed bats (Phyllostomidae) use three major senses, echolocation, olfaction and vision, for navigation and foraging (e.g., Bloss 1999; Laska 1990; Reiger et al. 1988; Schnitzler et al. 2003; Winter et al. 2003), which makes these organisms an exceptional system for investigating the relative role of different sensory modes during ecologically important tasks. Although there has been recent work on phyllostomid vision, including comparisons of their short- and long-wave opsins (Muller et al. 2007; Muller et al. 2009), the importance of vison in ripe fruit location and selection has not been experimentally tested in phyllostomids. By contrast, experimental evidence strongly suggests that omnivorous phyllostomids rely primarily on echolocation to locate fruits (Kalko and Condon 1998), and specialized frugivores employ either olfaction or a combination of olfaction and echolocation to locate ripe fruit (Hodgkison et al. 2007; Hodgkison et al. 2013; Kalko & Condon, 1998; Korine & Kalko 2005; Kalko & Schnitzler 1998).

The relative importance of different sensory modes for fruit detection may depend on which plant cues can be readily perceived within a specific environmental context. To date, it is unknown whether and how frugivorous phyllostomids integrate their primary sensory modes (echolocation and olfaction) conditional on which plant cues are present. While plant scents can travel over short or long distances (Riffell et al. 2014), they are rarely directional and may be difficult to detect in saturated scent environments such as rainforests. Conversely, echolocation allows for highly precise prey detection (Schnitzler and Kalko 2001; Brinkløv et al. 2009; Jakobsen et al. 2013), but phyllostomids emit short, high frequency and low intensity calls (Thies et al. 1998; Korine and Kalko 2005). The echoes from these calls provide information about size, shape, texture, range and position of an object in space relative to the bat (Simmons et al. 1975, 1983; Simmons and Stein 1980; Neuweiler 1989; Schmidt et al. 2000; Schnitzler and Kalko 2001), but for these types of information are only effective at very short distances because low intensity, high frequency calls attenuate rapidly in warm, humid environments (e.g., 45 −90 kHz attenuate at 1.4 to 4 dB/m at 25° C and 80% humidity; Jakobsen et al. 2013). Additionally, surrounding foliage can produce acoustic masking effects that may complicate fruit detection (Arlettaz et al. 2001, Korine and Kalko 2005). Therefore, flexibility in olfaction versus echolocation use could be highly beneficial for frugivorous bats given the limitations of each sensory mode within complex forest environments.

Here, we study the main plant cues and how this relates to the roles of echolocation and olfaction for fruit detection and localization in the chestnut short-tailed fruit bat, *Carollia castanea*, a highly abundant frugivore and ecologically important seed disperser that inhabits forests in Central and South America (Bonaccorso 1979; Fleming 1991) (Figure 1). This research builds upon the seminal work of Thies et al. (1998), which demonstrated the importance of olfaction for fruit detection in *Carollia;* here we investigate the relative contributions of vegetative and fruit scent cues that drive *Carollia’s* foraging decisions, and how varying foraging scenarios (i.e. changing of plant scent cues) may affect the relative reliance of these bats on olfaction versus echolocation for fruit detection. *Carollia castanea* feeds primarily on infructescences of Neotropical *Piper* plants (Piperales: Piperaceae), particularly *P. scintillans* (previously *P. sancti-felicis*; Hammel et al. 2003) in our study locality (Lopez and Vaughan 2007). We test if foraging decisions in *C. castanea* are primarily driven by clear signals of food availability (i.e., *P. scintillans* ripe fruit scent). We predict bats will cue in on ripe *P. scintillans* fruit scent and secondarily on its vegetation scent, since vegetation is a salient and fragrant part of the plant that could also aid in fruit localization. Since this prediction relies on the assumption that ripe fruit and vegetation scents differ in their chemical composition, we present analyses contrasting the volatile chemical composition of these plant parts. We also hypothesize that *C. castanea* potentially re-weight their sensory inputs to account for available environmental cues, and predict that foraging bats emit echolocation calls more frequently when scent cues are absent. This is because *Carollia* can use echolocation for the final localization of fruit at close range (Thies et al. 1998; Corlett 2011) and perhaps also when searching for potentially edible fruit patches at a longer range. To test our hypotheses, we conducted a series of experiments to mimic the sensory challenges fruit bats may encounter in nature, and quantified differences in the bats’ behavioral responses when exposed to different sensory cues. Our study contributes to the understanding of what chemical cues bats use for fruit selection, what contexts facilitate alternating between sensory modes, and the behavioral and sensory adaptations fruit bats have evolved for foraging.

**Fig. 1.**
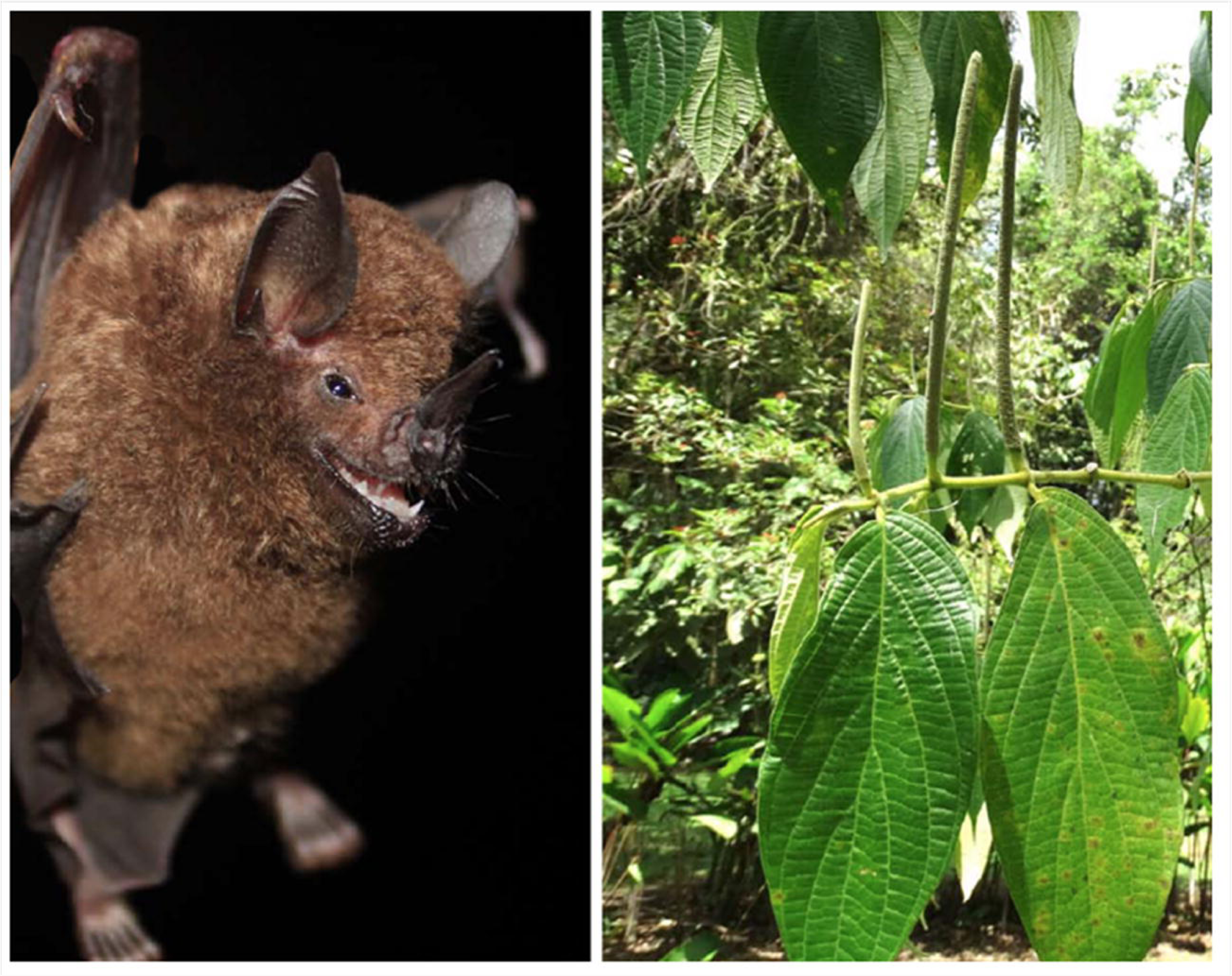
Study organisms, *Carollia castanea* (left) and *Piper scintillans* (right)

## MATERIAL AND METHODS

### Study animals

We used mist nets to capture *Carollia castanea* along forest trails at La Selva Biological Station in Sarapiquí, Heredia Province, Costa Rica (Supplementary Table 1). All individuals were experimentally naïve and were used in experiments on the night of capture. Upon capture, each bat was kept in a clean cotton bat bag prior to experiments. We conducted experiments on 21 bats (16 adult males and 5 adult non-lactating, non-pregnant females), and collected biometric data and a 2-3 mm wing biopsy (Disposable Biopsy Punches, Integra Miltex) from the uropatagium of each individual that had positive trials (see below). This was done for future genetic analyses, and helped ensure we did not use recaptured individuals in subsequent experiments. All individuals were released near the site of capture after the behavioral experiments and processing were completed. All procedures were approved by the University of Washington Institutional Animal Care and Use Committee (protocol# 4307-02).

**Table 1.**
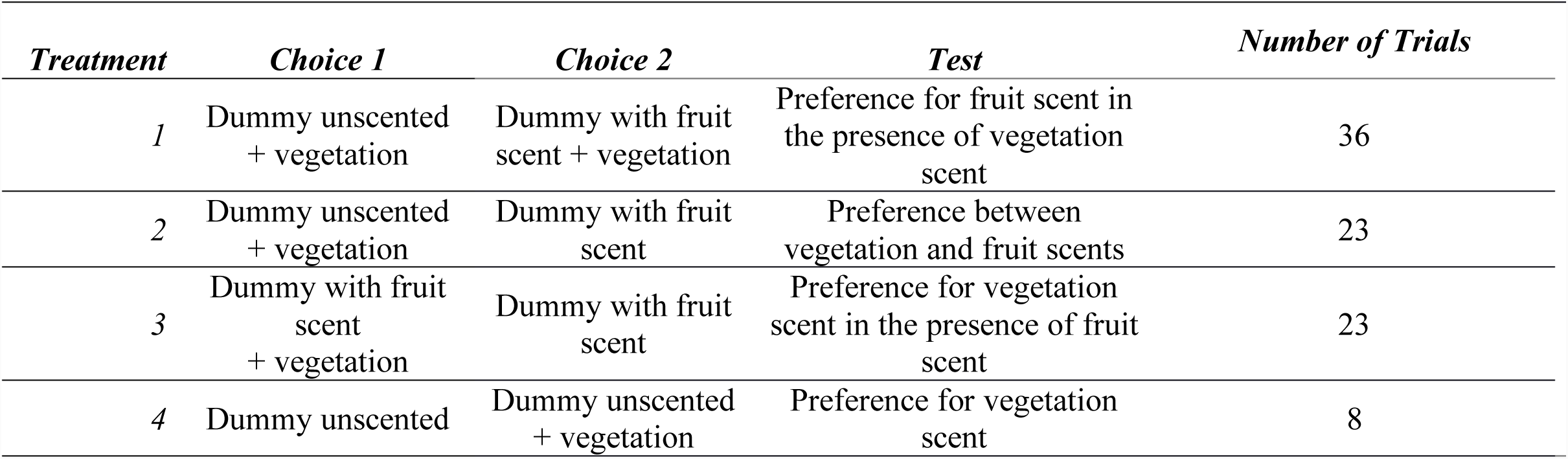
Description of the two target choices offered within each experimental treatment, and the response being tested during behavioral experiments on *Carollia castanea*. Number of trials differed among treatments due to differences in the number of positive responses of experimental bats (see Methods)

| *Treatment* | *Choice 1* | *Choice 2* | *Test* | *Number of Trials* |
| --- | --- | --- | --- | --- |
| 1 | Dummy unscented<br>+ vegetation | Dummy with fruit<br>scent + vegetation | Preference for fruit scent in<br>the presence of vegetation<br>scent | 36 |
| 2 | Dummy unscented<br>+ vegetation | Dummy with fruit<br>scent | Preference between<br>vegetation and fruit scents | 23 |
| 3 | Dummy with fruit<br>scent<br>+ vegetation | Dummy with fruit<br>scent | Preference for vegetation<br>scent in the presence of fruit<br>scent | 23 |
| 4 | Dummy unscented | Dummy unscented<br>+ vegetation | Preference for vegetation<br>scent | 8 |

### Experimental set-up

We conducted two-choice behavioral experiments without reward inside a flight cage (Coleman, 10 × 10 × 7 ft.) under natural ambient conditions at La Selva. As shown in Figure 2, we placed an infrared-sensitive handycam (4K HD Video Recording, Sony, Japan) on a tripod (30 cm from the ground), which allowed us to record the bats’ foraging behaviors under infrared light conditions (>700nm; beyond the spectral range of vision of phyllostomids; Jones et al. 2013). We recorded the bats’ echolocation calls with a condenser microphone (microphone capsule CM16, CMPA preamplifier unit, Avisoft Bioacoustics, Berlin, Germany) mounted at the top, center of the flight cage. During experiments, we visualized real-time calls using an ultrasound acquisition board (UltraSoudGate 116, Avisoft Bioacoustics, Germany; sampling rate 375 kHz, 16-bit resolution). At the back end of the flight cage, we placed a custom-built platform (90 cm long by 125 cm tall) which held two 50 ml falcon tubes, 40 cm apart, onto which we mounted each of the target (“fruit”) choices (Figure 2). To control for size and shape of these targets, we used 3D-printed dummy fruits (Form 2 printer with FGLPWH02 resin) of the same shape and size of an average *Piper scintillans* fruit. To mimic a ripe fruit, we smeared a dummy (3D printed) fruit with a standardized amount of ripe *P. scintillans* fruit pulp (approx. 0.62 g, ∼1/3 of a total fruit) collected on the same day of the experiments. We harvested vegetation (branches) from the same plants from which we collected ripe fruit. In trials with vegetation present, we placed the vegetation at the base of the dummy fruit, which is the natural configuration within the plant. Between each night of experiments, we cleaned dummy fruits with 95% ethanol to remove scents, rinsed them with water, and let them air dry at least 24 hours before reusing.

**Fig. 2.**
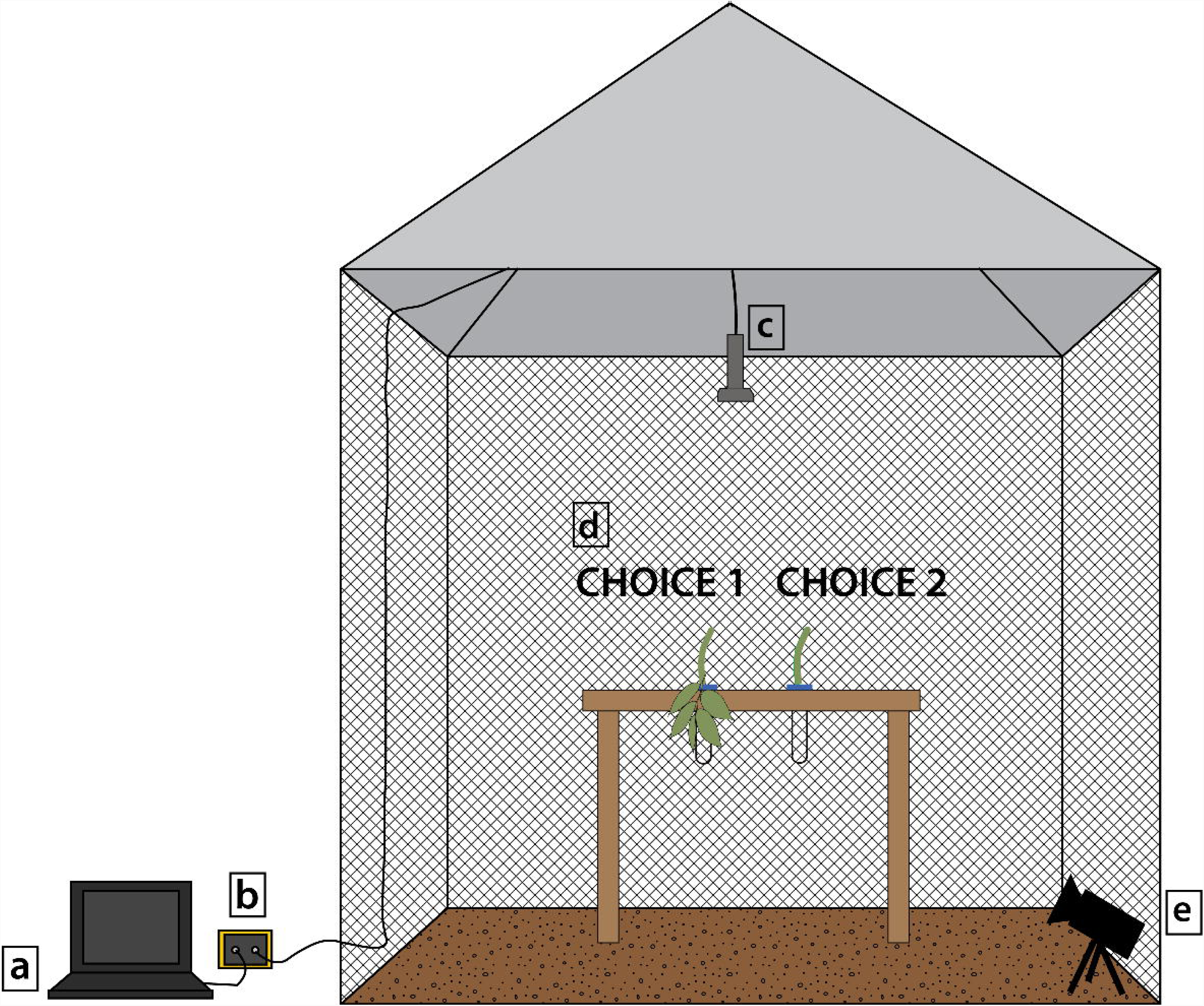
Diagram of experimental set-up: echolocation calls were visualized and recorded via a Dell 14 Rugged Extreme laptop (a) connected to a USG 116H recorder (b) that was connected to a CM16 condenser microphone (c). Target choice options were offered on a custom-made platform (d), here showing two example choice options, Choice 1: dummy with fruit scent and vegetation, and Choice 2: dummy with fruit scent only. Bat behaviors were recorded with a Sony infrared-sensitive handycam (e)

**Fig. 3.**
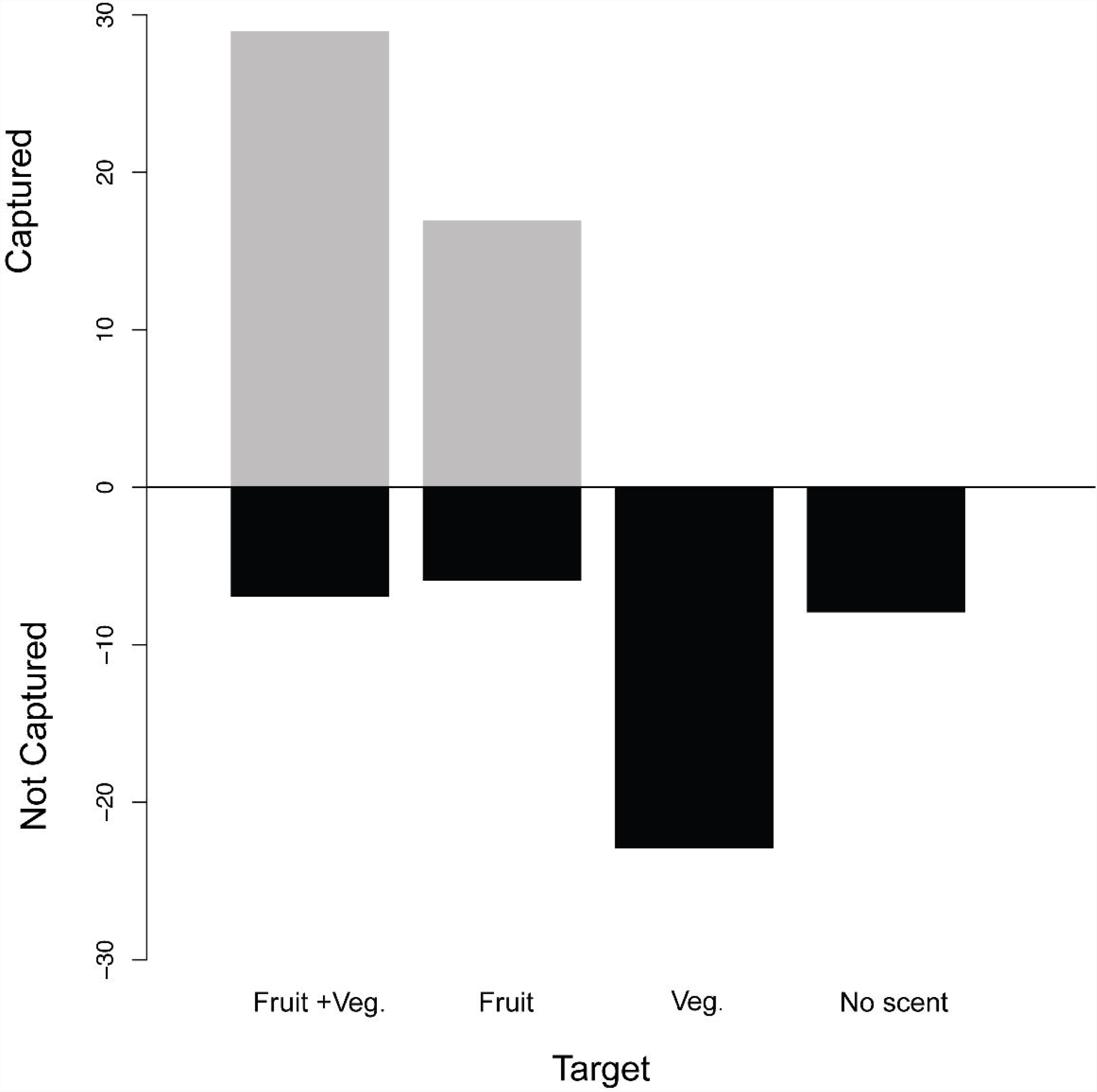
Summary of successful target captures by *Carollia castanea* for each of the four target types across behavioral experiments: dummy with fruit scent and vegetation (Fruit + Veg.), dummy with fruit scent only (Fruit), dummy unscented with vegetation only (Veg.), and dummy unscented (No scent)

To test our hypotheses, we presented each bat with a choice between two of the following targets during each experimental trial: 1) dummy with fruit scent (pulp from *P. scintillans*), 2) dummy with vegetation only, 3) dummy with fruit scent and vegetation, and 4) dummy with no fruit scent or vegetation (Table 1). We ran each trial for a maximum of 20 minutes per bat, and subjected bats to up to four trials, conditional on their performance on the initial trial. If the bat did not perform within 20 minutes, we released the individual. Conversely, if the bat attempted to capture the fruit within the 20-minute duration of the trial, we considered it a positive trial and began a new trial. To begin a new trial, one of us entered the flight cage and switched out choice targets. We randomized both the order we presented each trial and the position (left, right) of the target choice on the platform between consecutive trials to minimize confounding effects due to bat spatial learning (Thiele and Winter 2004). At the end of trials, we used a hand net to recapture the bat inside the flight cage, processed and released it, as described above.

### Analysis of flight behavior during target search and approach

We watched videos of the behavioral trials at normal playback speed on a computer at the University of Washington, Seattle, WA. From each video, we recorded: the amount of time it took the bat to capture one of the target choices presented, the real time of capture (to synchronize with acoustic calls), and the individual’s choice. We defined a capture event as a bat landing on a target choice and attempting to bite it. We noted additional characteristics of the bat’s flight behavior (e.g., exploratory flights around the flight cage) and target exploration (i.e., hovering over fruit) for all trial videos.

### Analysis of echolocation behavior during target approach

We analyzed echolocation calls emitted during target approach using Avisoft SASLabPro v. 4.40 (Avisoft Bioacoustics, Berlin, Germany). We used the time of the capture events from video recordings and matched them with the time stamps of the call files to synchronize acoustics with recorded capture events. These files were used in the subsequent analyses and included the acoustic calls for one minute prior to the capture event, which we defined as the search/approach window. We chose a minute interval prior to capture because we were not only interested in sensory behavior for target localization (typically seen in the approach phase), but also the sensory behavior when ‘searching’ for food. *Carollia castanea*, similar to other phyllostomids, emits calls well above 20kHz (Thies et al. 1998; Brinkløv et al. 2011), so we used this as the cut-off frequency to avoid including noise from recording at high gain in ambient conditions (Geipel et al. 2013). We filtered each acoustic sequence using a high-pass filter (at 20 kHz) and visualized spectrograms using a Hamming window (512 fast Fourier transform, 98.95% overlap). We extracted the following echolocation call parameters for comparisons across trial types: maximum frequency (kHz), minimum frequency (kHz), peak frequency (i.e. frequency with the highest amplitude, kHz), call duration (ms), call interval (ms), and total bandwidth (kHz) from the spectrograms at the maximum energy of each call. We compiled sequences per individual (approximately 8 – 20) and calculated mean and standard deviation for each call parameter per trial type per individual (Table 2).

**Table 2.** Means (± standard deviation) of echolocation call parameters for *Carollia castanea* during each experimental treatment (from Table 1). Means for the call parameters were calculated by averaging the calls for each individual bat across its entire calls sequence (the approach call for one individual, for one treatment), and averaging each of these values across individuals within each treatment

| Treatment | Duration (ms) | Interval (ms) | Peak Frequency (kHz) | Min. Frequency (kHz) | Max. Frequency (kHz) | Bandwidth (kHz) |
| --- | --- | --- | --- | --- | --- | --- |
| 1 | $2.09 \pm 1.02$ | $314.7 \pm 252.8$ | $83.9 \pm 3.44$ | $62.5 \pm 10.0$ | $108.3 \pm 9.131$ | $45.8 \pm 15.0$ |
| 2 | $1.34 \pm 0.0103$ | $268.1 \pm 151.5$ | $85.9 \pm 6.89$ | $68.0 \pm 9.79$ | $111.7 \pm 12.54$ | $43.7 \pm 19.1$ |
| 3 | $1.70 \pm 0.0589$ | $421.8 \pm 20.63$ | $83.1 \pm 5.69$ | $60.7 \pm 6.16$ | $111.3 \pm 4.531$ | $50.6 \pm 4.92$ |
| 4 | $2.71 \pm 0.206$ | $81.6 \pm 26.8$ | $87.2 \pm 10.65$ | $64.7 \pm 10.7$ | $108.9 \pm 11.51$ | $44.2 \pm 11.6$ |

### Statistical analyses of behavioral experiments

We performed all statistical analyses in R v. 3.2.4 (R Core Team, 2018). We used chi-squared tests to assess the differences in bat preference among target types. We tested the normality of the echolocation call data using Shapiro-Wilks tests (Shapiro and Wilk 1965) and subsequently log-transformed these data to improve normality. We compared the differences in call parameters among trial types using analysis of variance (ANOVA).

### Target comparisons

To determine if *P. scintillans* ripe fruit and vegetation present different olfactory cues to *C. castanea*, we compared the volatile organic compounds (VOCs) that make up the scents of these plant parts. We collected vegetation (7.0 – 7.8 g of leaf material; one branch) and ripe fruit (19.9 –20.6 g; 13 – 15 fruits) samples from four *P. scintillans* plants at La Selva. This larger sample of ripe fruit was necessary for VOC capture and detectability by our experimental setup (below). We collected VOCs from these samples via dynamic headspace adsorption using a push–pull system (Raguso and Pellmyr, 1998; Riffell et al., 2008). Within two hours of collection, we placed each sample in a 3L teflon bag (Reynolds, Richmond, VA, USA) and connected the bag to a diaphragm pump (400-1901, Barnant Co., Barrington, IL, USA) that pulled the fragrant headspace air through a sorbent cartridge trap (50 mg Porapak Q with silanized glass wool Waters Corp., Milford, MA, USA) and pushed air though a charcoal filter. We collected VOCs in this manner for 20 hours per sample, following previously established protocols and to ensure characterization of the full chemical profile (Byers et al. 2014). We eluted trapped volatiles from each sample’s sorbent cartridge with 600 µl of HPLC-grade hexane into a 2mL borosilicate glass vial with a Teflon-lined cap. Subsequently, we stored all of the samples at –20°C to –80°C. We analyzed a 3 µl aliquot of each sample using an Agilent 7890A GC (gas chromatograph) and a 5975C Network Mass Selective Detector (Agilent Technologies, Palo Alto, CA, USA). To separate the VOCs, we used a DB-5MS GC column (J&W Scientific, Folsom, CA, USA; 30 m, 0.25 mm, 0.25 µm) with helium as the carrier gas flowing at a constant rate of 1 cc per min (Byers et al. 2014). The initial oven temperature was 45°C for 4 min, followed by a heating gradient of 10°C min^−1^to 230°C, which was then held isothermally for 4 min. We initially identified the chromatogram peaks with the aid of NIST 08 mass spectral library (v. 2.0f; ca. 220,460 spectra of 192,108 different chemical compounds) followed by verification using alkane standards and comparing with published Kovats indices. We integrated the peaks for each compound using ChemStation software (Agilent Technologies) and present them in Table 4.

## RESULTS

### Target preference

Most individuals performed exploratory flights prior to showing interest in the presented target choices. These behaviors consisted of circling flights around the cage without approaching the target. In most trials (83%), bats attempted to capture a target by landing and trying to bite the dummy fruits (Supplementary Video 1). Comparisons across treatments revealed that *C. castanea* strongly preferred targets consisting of a dummy fruit with fruit scent (n = 10, χ^2^= 6.21, P = 5.69e-05) or a dummy fruit with fruit scent and vegetation (n = 13, χ^2^= 22.154, P = 2.52e-06) over targets that had an unscented dummy fruit and vegetation. The presence of vegetation did not affect the bats’ preferences for fruit scent (dummy fruit with fruit scent vs. dummy fruit with fruit scent and vegetation: n = 13, χ^2^= 0, P = 0.99). Bats never chose unscented dummy fruits, either alone or with vegetation.

### Echolocation behavior

All bats used in the experiments emitted echolocation calls throughout the trials. We did not find statistically significant differences in the echolocation call parameters among treatment types during the search/approach window (minimum frequency, maximum frequency, peak frequency, bandwidth, duration, pulse interval, all P > 0.05; Table 3). However, there were marked trends in call duration and interval in some trial types. Bats emitted echolocation calls more frequently (shorter interval) and of longer duration in treatments where no fruit scent was present (unscented dummy vs. unscented dummy with vegetation; Figure 4).

**Table 3.** Summary of ANOVA results comparing each call parameter trait across the four experimental treatments (from Table 1).

| Variable | Df | Sum. Sq. | Mean Sq. | F-Value | P(>F) |
| --- | --- | --- | --- | --- | --- |
| Duration | 1 | 0.852 | 0.852 | 0.728 | 0.405 |
| Interval | 1 | 1.37 | 1.37 | 1.209 | 0.287 |
| Peak Freq. | 1 | 0.36 | 0.361 | 0.194 | 0.665 |
| Min. Freq. | 1 | 0.00 | 0.0001 | 0.00 | 0.995 |
| Max. Freq. | 1 | 0.00 | 0.002 | 0.001 | 0.975 |
| Bandwidth | 1 | 0.00 | 0.0004 | 0.000 | 0.989 |

**Table 4.** Volatile organic compounds found in the scent of *Piper scintillans* ripe fruit and vegetation. Values are the mean proportion across all samples (n = 2 fruit; n = 2 vegetation). KRI: Kovats retention indices; IUPAC: International Union of Pure and Applied Chemistry nomenclature

| Chemical Name | IUPAC | KRI | Class | Ripe fruit | Vegetation |
| --- | --- | --- | --- | --- | --- |
| 3-Hexene-1-ol | (Z)-hex-3-en-1-ol | 858 | aliphatic alcohol | 0 | 0.343 |
| β-Pinene | 6,6-dimethyl-4-methylidenebicyclo[3.1.1]heptane | 981 | monoterpene | 0.089 | 0.067 |
| α-Phellandrene | 2-methyl-5-propan-2-ylcyclohexa-1,3-diene | 1008 | monoterpene | 0.01 | 0 |
| 3-Carene | 4,7,7-trimethylbicyclo[4.1.0]hept-3-ene | 1010 | monoterpene | 0.034 | 0 |
| α-Terpinene | 1-methyl-4-propan-2-ylcyclohexa-1,3-diene | 1020 | monoterpene | 0.015 | 0.053 |
| p-Cymene | 1-methyl-4-propan-2-ylbenzene | 1028 | aromatic | 0.08 | 0.072 |
| β-Ocimene | 3,7-dimethylocta-1,3,6-triene | 1050 | monoterpene | 0.033 | 0 |
| γ-Terpinene | 1-methyl-4-propan-2-ylcyclohexa-1,4-diene | 1064 | monoterpene | 0.018 | 0.055 |
| Terpinolene | 1-methyl-4-propan-2-ylidenecyclohexene | 1091 | monoterpene | 0 | 0.023 |
| Ethyl benzoate | ethyl benzoate | 1175 | aromatic | 0.071 | 0.126 |
| α-Cubebene | 4,10-dimethyl-7-(propan-2-yl)tricyclo[4.4.0.0<sup>1,5</sup>]dec-3-ene | 1354 | sesquiterpene | 0.022 | 0 |
| β-Elemene | 2,4-Diisopropenyl-1-methyl-1-vinylcyclohexan | 1394 | sesquiterpene | 0.178 | 0.06 |
| α-Cedrene | Cedr-8-ene | 1419 | sesquiterpene | 0.023 | 0.024 |
| β-Caryophyllene | 4,11,11-trimethyl-8-methylidenebicyclo[7.2.0]undec-4-ene | 1433 | sesquiterpene | 0.208 | 0.036 |
| α-Bergamotene | 4,6-dimethyl-6-(4-methylpent-3-enyl)bicyclo[3.1.1]hept-3-ene | 1440 | sesquiterpene | 0.016 | 0.032 |
| α-Caryophyllene | 2,6,6,9-tetramethylcycloundeca-1,4,8-triene | 1470 | sesquiterpene | 0.029 | 0.011 |
| γ-Muurolene | 7-methyl-4-methylidene-1-propan-2-yl-2,3,4a,5,6,8a-hexahydro-1H-naphthalene | 1485 | sesquiterpene | 0 | 0.021 |
| Germacrene D | 1-methyl-5-methylidene-8-propan-2-ylcyclodeca-1,6-diene | 1495 | sesquiterpene | 0.157 | 0.034 |
| Alloaromadendrene | 1,1,7-trimethyl-4-methylidene-2,3,4a,5,6,7,7a,7b-octahydro-1aH-cyclopropa[e]azulene | 1504 | sesquiterpene | 0.011 | 0.028 |
| Caryophyllene oxide | 4,12,12-Trimethyl-9-methylene-5-oxatricyclo[8.2.0.0<sup>4,6</sup>]dodecane | 1581 | sesquiterpene | 0 | 0.009 |

**Fig. 4.**
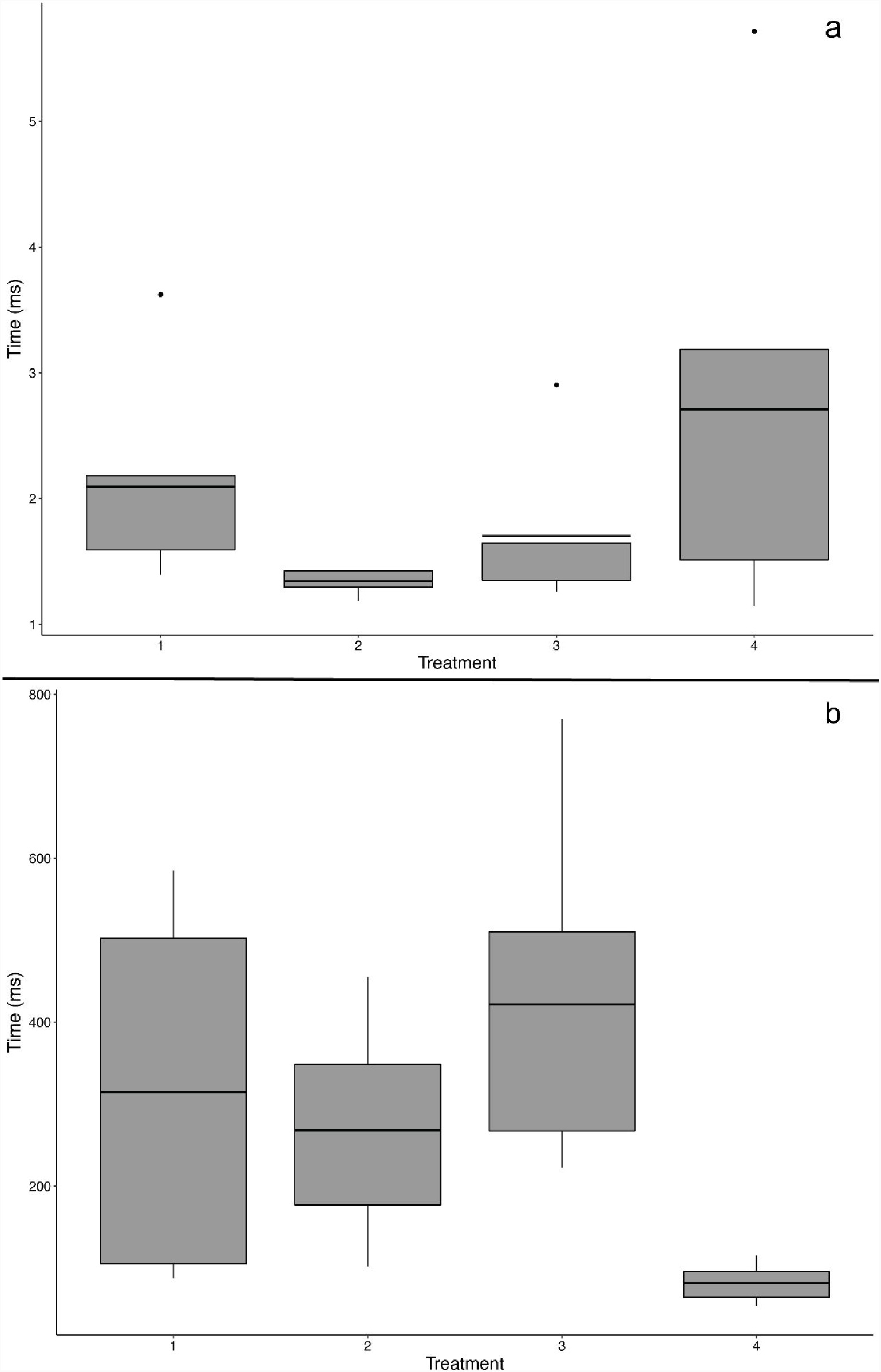
Summary of the duration of *Carollia castanea’s* echolocation calls across treatments (a), and summary of the interval of *Carollia castanea’s* echolocation calls across treatments (b). Treatments are described in Table 1

### Chemical differences between targets

The scent profiles of *P. scintillans* vegetation and ripe fruit differ slightly in VOC composition, and greatly in the proportion of specific VOCs (Table 4). The vegetation scent of *P. scintillans* is dominated by 3-hexene-1-ol, which is not found in the ripe fruit scent. Conversely, the ripe fruit scent is characterized by a greater abundance of *β*-caryophyllene, germacrene D and *β*-elemene. Both vegetation and fruit scents have a low abundance of p-cymene and *β*-pinene (Table 4).

## DISCUSSION

An animal’s sensory ecology and behavior often reflects the environment it inhabits, as well as its evolutionary history. As such, certain sensory modalities play key roles in mediating ecological interactions. Mammalian frugivores are able to locate and acquire ripe fruit by using and integrating across sensory modalities: they use vision to detect differences in fruit color and luminance (Burkhardt 1989; Osorio and Vorobyev 2008; Melin et al. 2008; Hiramatsu et al. 2008; Valenta et al. 2013), olfaction to detect individual volatile organic compounds (VOCs) or entire odor plumes (Sánchez et al. 2006; Valenta et al. 2013) and, in the case of phyllostomid bats, echolocation to gather information about fruit shape and location (Helversen & Helversen 1998; Kalko & Condon 1998). Our results indicate that *C. castanea* makes foraging decisions based on ripe fruit scent over all other cues presented, but may rely more heavily on echolocation when olfactory cues are absent. Importantly, in the absence of plant scent cues, a *P. scintillans* fruit shape does not elicit a prey capture response from *C. castanea*. This gives insight to the primary role of olfaction, followed by echolocation, when these bats forage for fruit.

The importance of olfaction for foraging in *C. castanea* is supported by previous research on other *Carollia* species that rely less on *Piper* as a food resource. For example, *C. perspicillata* can recognize minute concentrations of particular chemical components (fruit-typical odor components like ethyl butyrate, n-pentyl acetate, or linalool; Laska 1990), are attracted to fruit scent even when no other cues are present (Hessel and Schmidt 1994), and visit mist nets spiked with the essential oil of *Piper gaudichaudianum* more frequently (Mikich et al. 2003). Here, we link the bouquet of VOCs from a known, preferred food source with the behavioral preferences of *C. castanea*. Our experimental results strongly suggest that *C. castanea* uses ripe fruit scent, as opposed to a combination of ripe fruit and/or vegetation scent, or fruit shape, as the cue to locate food items. Our chemical analyses of ripe fruit and vegetation VOCs provide an explanation for this pattern: the scent profile of the *P. scintillans* ripe fruit and vegetation are somewhat similar in composition, but differ greatly in abundance of some specific VOCs. Additionally, the ripe fruit scent profile contained a few distinct VOCs that were not found in vegetation, and vice versa. Considering that *C. castanea* forages in a complex sensory environment, the forest understory, it may be advantageous for the bats to cue in specific chemicals that unmistakably signal fruit ripeness against the background of unripe fruit and vegetation within a *Piper* bush, as well as adjacent vegetation. Our results motivate future work to examine whether some of these key volatiles or their ratios may signal fruit ripeness amongst the vegetative mélange.

Echolocation call parameters did not differ significantly in frequency between trial types, suggesting that *C. castanea* has a stereotyped call structure regardless of their foraging tasks. While this has not been broadly studied, having a stereotyped echolocation call is common in phyllostomids (Kalko and Condon 1998; Thies et al. 1998; Korine and Kalko 2005; Geipel et al. 2013). Nevertheless, our experiments revealed that *C. castanea* potentially modulates time-linked echolocation traits (i.e., duration of the call and time between calls, interval) when confronted with different prey cues. As in all mammals, phyllostomid bats process olfactory cues by inhaling air through their nose, but also emit echolocation calls out of their nose. Because of the potential conflict between these two functions performed by the nasal cavity, we propose that frugivorous phyllostomids exhibit behavioral modulation in their nasally-linked senses to alternate between and maximize effectiveness of one sensory cue versus another, when appropriate.

Echolocation provides bats with high-resolution information about shape, surface texture, and material of an object at close range (Schnitzler et al. 1983; Ostwald et al. 1988; Kober and Schnitzler 1990), but bats also use echolocation for navigation and detection of plants that signal through morphology for better acoustic detection. We saw a general trend of longer duration of echolocation calls and shorter intervals between calls when bats were offered choices that did not include a ripe fruit scent cue. Decreased time between calls (interval) and longer duration means these bats were calling more frequently in the absence of ripe fruit scent. We hypothesize that, when ripe fruit odor cues are absent, *C. castanea* relies more heavily on echolocation to locate a potential food item, in this case one that may resemble an edible *Piper* fruit. In contrast, when bats were presented with any target that had ripe fruit scent (one or two choices), they emitted shorter echolocation calls at longer intervals, thus echolocating less frequently. We hypothesize that the decrease in echolocation call duration and an increase in interval could be linked to an increase in the bats’ use of olfaction as they attempt to locate edible ripe fruits or determine which one is the ‘most edible’ option.

If ripe fruit scent is the primary cue for fruit location and selection by *C. castanea*, why are there differences in echolocation call duration and interval between treatments with and without ripe fruit scent? Previous studies have demonstrated that phyllostomids bats can use echolocation to determine the position of a fruit (Kalko and Condon 1998), and echolocating bat species, in general, alter call parameters to overcome acoustic masking effects during prey location (Kalko and Schnitzler 1993). This can be accomplished by changing the duration and interval of the call (Siemers and Schnitzler 2000). Bats typically extend the duration of a call when searching for prey and during orientation flights (Kalko and Schnitzler 1993), and decrease the time between calls when approaching a prey item (Kalko and Schnitzler 1993; Siemers and Schnitzler 2000). In our experiments, bats never chose unscented dummy fruits as potential food options, but our behavioral and acoustic recordings demonstrated that they did explore them via echolocation. We propose that *C. castanea* has a series of criteria (e.g., fruit scent, shape, configuration of fruit in relation to vegetation), which may be hierarchical, and are integrated during the search and localization of a potential food item.

The use of echolocation and olfaction for food selection has been documented in other frugivorous and omnivorous phyllostomids. *Artibeus jamaicensis* is a specialized frugivore that detects, localizes and classifies ripe fruits primarily by olfaction (Kalko et al. 2010). In contrast, *Phyllostomus hastatus*, a large omnivorous bat, consistently uses echolocation over olfaction when foraging for *Gurania spinulosa*, a pendulous fruit-bearing vine (Kalko and Condon 1998). These examples illustrate an echolocation-olfaction continuum across phyllostomids that forage for fruit, and suggests that multiple sensory modes are important for fruit foraging in complex environments. They also substantiate that sensing mode could be conditional on which food cues are present or the degree of specialization of each species (e.g., omnivores vs. specialized frugivores). To date, it is still unclear which senses are most important for fruit foraging in most bat frugivores, and what facilitates the use of one sense over another.

This study provides behavioral links between a frugivore’s sensory abilities and plant cues, a relationship that is critical to understanding the ecological dynamics and coevolution between plants and their seed dispersers. There is still much to learn about how vertebrate frugivores perceive and interact with their potential food sources, thus further observational and experimental studies are critical for determining what specific fruit traits (e.g., compounds or combination of compounds in ripe fruit) drive fruit selection by frugivorous species.

All procedures performed in studies involving animals were approved by and in accordance with the ethical standards of the University of Washington Institutional Animal Care and Use Committee (protocol# 4307-02)

## Data Availability

The datasets generated during and analyzed during the current study are available from the corresponding author on reasonable request.

## Acknowledgements

We would like to thank Dr. Stephanie Smith for contributions in field work and David Villalobos for help with permits. We would also like to thank the staff at La Selva Biological Research Station, especially Orlando Vargas Ramírez, Bernal Matarrita Carranza and Danilo Brenes Madrigal for their invaluable help during fieldwork. This work was supported by the National Science Foundation (grant number 1456375 to SES and JAR), and a Tinker Field Research Grant (to MEL).

