## Supplemental Table 1 for "A fruitful endeavor: scent cues and echolocation behavior used by *Carollia castanea* to find fruit"

**Supplementary Table 1** List of individual bats used in experiments and associated biometric data: M(male), F(female), A (adult), NP-NL (Not pregnant, not lactating), NS (non-scrotal)

| No. | Species | ID | Sex | Age | Rep. Cond. | Mass (g) | FA (mm) |
|---|---|---|---|---|---|---|---|
| **1** | Carollia castanea | 083016_1 | F | A | NP-NL | 14 | 37.3 |
| **2** | Carollia castanea | 083016_2 | M | A | NS | 12 | 35.8 |
| **3** | Carollia castanea | 083016_3 | M | A | NS | 11 | 37.4 |
| **4** | Carollia castanea | 083116_1 | M | A | NS | 14 | 37.9 |
| **5** | Carollia castanea | 083116_2 | M | A | NS | 12 | 36.1 |
| **6** | Carollia castanea | 083116_3 | F | A | NP-NL | 14 | 37.1 |
| **7** | Carollia castanea | 090116_2 | M | A | NS | 13 | 36.8 |
| **8** | Carollia castanea | 090116_3 | M | A | NS | 14 | 37.6 |
| **9** | Carollia castanea | 090816_2c | M | A | NS | 14 | 36.7 |
| **10** | Carollia castanea | 090816_3c | M | A | NS | 13 | 36.3 |
| **11** | Carollia castanea | 090816_4c | F | A | NP-NL | 14 | 36.5 |
| **12** | Carollia castanea | 090816_5c | M | A | NS | 11 | 36 |
| **13** | Carollia castanea | 090816_6c | M | A | NS | 11 | 36.2 |
| **14** | Carollia castanea | 090916_1 | F | A | NP-NL | 14 | 38.7 |
| **15** | Carollia castanea | 090916_2 | M | A | NS | 14 | 37.1 |
| **16** | Carollia castanea | 090916_3 | M | A | NS | 11 | 37.2 |
| **17** | Carollia castanea | 090916_5 | M | A | NS | 12 | 36.2 |
| **18** | Carollia castanea | 091016_1 | F | A | NP-NL | 12 | 36.2 |
| **19** | Carollia castanea | 091016_2 | M | A | NS | 13 | 37.2 |
| **20** | Carollia castanea | 091016_4 | M | A | NS | 14 | 37.1 |
| **21** | Carollia castanea | 091216_3 | M | A | NS | 10 | 36.15 |
